# Comparison of dsDNA and ssDNA-based NGS library construction methods for targeted genome and methylation profiling of cfDNA

**DOI:** 10.1101/2022.01.12.475986

**Authors:** Jianchao Zheng, Zhilong Li, Xiuqing Zhang, Hongyun Zhang, Shida Zhu, Jianlong Sun, Yuying Wang

## Abstract

Cell-free DNA (cfDNA) profiling by next generation sequencing (NGS) has wide applications in cancer diagnosis, prognosis, and therapy response monitoring. One key step of cfDNA deep sequencing workflow is NGS library construction, whose efficiency determines effective sequencing depth, sequencing quality, and accuracy. In this study, we compared two different cfDNA library construction methods for the applications of mutation detection and methylation profiling: the conventional method which captures double-stranded DNA (dsDNA) molecules, namely the dsLib workflow, and an alternative method which captures single-stranded DNA (ssDNA), namely the ssLib workflow. Our results suggest that the dsLib method was preferrable for mutation detection while the ssLib method proved more efficient for methylation analysis. Our findings could help researchers choose more appropriate library construction method for corresponding downstream sequencing applications.

## Introduction

Cell-free DNA (cfDNA), primarily derived from cell apoptosis, has been shown to be an important biomarker of many physiological and pathological conditions such as autoimmunity, infection, pregnancy, exercise, transplantation, and cancer [1-3]. It is detectable in almost all body fluids including plasma, serum, and urine, with a peak size at approximately 166 bp [4, 5]. In cancer patients, there is a subset of cfDNA known as circulating tumor DNA (ctDNA), which originates from tumor cells and carries genetic and epigenetic characteristics of the tumor [6, 7]. cfDNA has a short half-life in circulation (reported to be between 16 and 150 minutes), can be repeatedly sampled, and may potentially overcome intratumor heterogeneity compared to tissue biopsies [8]. These unique characteristics make cfDNA-based liquid biopsy an ideal approach for cancer diagnosis, prognosis, and therapy response monitoring [9, 10]. However, due to low concentration of cfDNA in plasma (∼3 ng/ml in healthy individuals) and very small fraction of ctDNA among abundant cfDNA that derived from blood cells and normal tissues, accurate detection of ctDNA remains a challenging task [11, 12].

Massively parallel sequencing (MPS), also known as next generation sequencing (NGS), has been widely applied in both research and diagnostic fields [13, 14]. Millions of DNA fragments can be simultaneously sequenced and analyzed by NGS [15]. Furthermore, targeted capture sequencing allows for deeper sequencing for target regions of interest at a lower cost [16]. Remarkably, efficient library construction before targeted capture sequencing determines effective sequencing depth and remains indispensable to successful sequencing of the target regions, and is particularly critical for sequencing of limited amount of cfDNA, and for identification of variants with lower allele fractions [9, 17]. During library construction, platform-specific adapters, which contain sample barcode sequence(s) and common primer binding sites for subsequent amplification and sequencing, are ligated to both ends of the original DNA fragments [18]. Various library construction methods have been introduced, aiming to improve DNA conversion efficiency (defined as the fraction of original DNA molecules that are successfully converted to the final library) [19-23]. Conventional double-stranded library (dsLib) construction workflow (such as what is used in the KAPA Hyper Prep kit) consists of following steps: (i) end repair and dA-tailing of the double-stranded (dsDNA) templates; (ii) adapter ligation; (iii) library amplification and purification [24]. On the other hand, single-stranded library (ssLib) construction was usually initialized by adapter ligation to single-stranded DNA (ssDNA) templates and followed by library amplification and purification [19, 22, 25]. The ssLib construction method was originally developed to recover ancient and/or degraded DNA fragments which are usually poorly captured by conventional dsDNA-based library preparation [26, 27]. Previously, researchers have compared their performance in applications such as non-invasive prenatal testing (NIPT), which is based on shallow whole-genome sequencing (WGS), and found no advantage for ssDNA-based methodology [28]. However, there hasn’t been systematic study to compare the performance of these two methods when used for cfDNA sequencing for cancer-related applications.

In this study, we compared the dsLib workflow and ssLib workflow for targeted deep sequencing (for variant detection) and methylation sequencing (for detection of cytosine methylation, an important form of epigenomic modification) of cfDNA, two applications important for cancer diagnosis. We found that, for targeted deep sequencing, the dsLib method achieved overall better performance and satisfactory limit of detection (LOD). For methylation sequencing, we compared the dsLib and ssLib workflow coupled with either bisulfite-based or enzyme-based cytosine conversion methods, and found that ssLib coupled with bisulfite conversion showed notably better performance.

## Materials and Methods

### Ethical Compliance

This study was approved by the institutional review board of BGI (NO. BGI-IRB: 19077).

### Sample collection and cfDNA isolation

After obtaining informed consent, blood samples were collected from 37 healthy volunteers and 2 lung cancer patients in 10 mL K2 EDTA BD Vacutainer tubes. Blood was separated immediately by an initial centrifugation at 1,600 × g for 10 min and then by a second centrifugation at 16,000 × g for 10 min. Plasma were pooled and split into 4ml per reaction for cfDNA isolation using MagPure Circulating DNA Maxi Kit (Magen, China) per manufacturer’s instruction. Extracted cfDNA samples from healthy volunteers were pooled together to obtain sufficient homogeneous material for subsequent analysis. cfDNA was quantitated by Qubit dsDNA High Sensitivity Kit (Thermo Fisher Scientific, USA). The 1% Multiplex I cfDNA reference standards HD778 (Horizon Discovery, UK) were spiked into healthy donor cfDNA at 0.1%, 0.25%, or 0.5% to simulate cfDNA samples with defined mutant allele frequencies (MAFs). Experiments were performed in triplicates.

### Double-stranded cfDNA library construction

Duplex unique molecular identifier (UMI) adapters for MGISEQ-2000 sequencer were designed according to principles described by Newman et al [21] with the modification that 3-bp UMIs were chosen instead of 2-bp UMIs in order to accommodate a higher library complexity. To avoid potential issues during sequencing caused by low complexity at the T-A ligation position (constant base), 32 pairs of UMI adapters were incorporated with an additional base (G or C) before the T-A ligation position. Long oligonucleotides UMIxxL (5’-Phosphorylation-[C/G/-]-NNNAAGTCGGAGGCCAAGCGGTCTTAGGAAGACAA -3’) and short oligonucleotides UMIxxS (5’-GACATGGCTACGATCCGACTNNN-[G/C/-]-T-3’) were synthesized by BGI tech solutions (Beijing Liuhe co.limited). Each oligo was dissolved to 100 μM using TE buffer. For each pair of adapters, 5 μL UMIxxL and 5 μL UMIxxS oligos (100 μM) were combined and brought up to 20 μL with TE buffer. Oligos were annealed for more than 30 minutes at room temperature. 64 UMI adapters (25 μM) were mixed and diluted to 5 μM, marked as UMI64M.

Double-stranded cfDNA libraries were prepared either by KAPA Hyper Prep kit (Kapa Biosystems, cat. No. KK8504) per manufacturer’s instruction or our custom library construction protocol. For the latter, briefly, 1-10 ng cfDNA was mixed with end-repair master mix consisting of T4 DNA polymerase (Enzymatics, cat. No. P7080L), T4 polynucleotide kinase (Enzymatics, cat. No. Y9040L), rtaq DNA polymerase (MGI, cat. No. 01E012MM), dNTP, and T4 DNA ligase buffer, and kept at 20□ for 30 min followed by 65□ for 30 min. Then UMI64M adapter was added to the end-repair reaction product and mixed by pipetting, followed by adding ligation master mix consisting of golden T4 DNA ligase (MGI, cat. No. 02E004MM), 10× T4 DNA ligase buffer, and PEG6000 (Sigma Aldrich, 50%). The ligation reaction was incubated at 16□ for 60 min. Adapter ligated DNA was purified using Agencourt AMPure XP beads. Next, index PCR was then performed and purified using Agencourt AMPure XP beads. The concentration of final library was determined by Qubit dsDNA High Sensitivity Kit.

Double-stranded cfDNA methylation sequencing libraries were prepared according to above library preparation workflow with following modifications: (i) 0.05 ng fragmented lambda DNA was spiked into the 10 ng cfDNA to monitor bisulfite conversion rate; (ii) all cytosines of adapter were methylated; (iii) after purification of the ligation product, bisulfite conversion was performed using EZ DNA Methylation Gold kit (Zymo Research, cat. No. D5006) or EM-seq Conversion Module (NEB, cat. No. E7125); (iv) index PCR was performed by 2×Golden U+ High-fidelity Readymix (MGI, cat. No. 01K01701MM).

### Single-stranded cfDNA library construction

The single-stranded library preparation method was based on the ssDNA2.0 method [19] with the modification that T-A ligation was used to further improve ligation efficiency. Briefly, MyOne C1 beads carrying the extension product were resuspended in the A-tailing reaction mix consisting of Klenow (3’-5’ exo-) (Enzymatics, cat. No. P7010-LC-L), 10× blue buffer, and dATP, and incubated at 37°C for 30 min then at 75°C for another 30min. The libraries were amplified by a specific number of PCR cycles based on cfDNA input amount, purified by Agencourt AMPure XP beads, and eluted in nuclease-free water.

Single-stranded cfDNA methylation sequencing libraries were prepared as above after input cfDNA was converted using either the EZ DNA Methylation Gold kit (Zymo Research, cat. No. D5006) or the EM-seq Conversion Module (NEB, cat. No. E7125). To monitor the conversion rate, 0.05 ng fragmented lambda DNA was spiked into 10 ng cfDNA.

### Target capture and sequencing

A custom capture panel that spans 220 kb and covers 139 cancer driver genes was designed and synthesized by IDT technologies as previously described [29]. Targeted genome capture was performed using xGen® Lockdown® Reagents (IDT technologies) and BGI adapter-specific blockers (BGI). 6 or 8 Libraries were pooled (400ng each) and captured per manufacturer’s instruction.

Targeted methylation capture was performed using a custom-designed 198kb panel of TargetCap methylation probes and reagents (BoKe Bioscience China, cat. No. MP121CD) and BGI adapter-specific blockers (BGI). 6 or 8 Libraries were pooled (400ng each) and captured per manufacturer’s instruction.

The above captured cfDNA genome or methylation libraries were amplified and purified with AMPure XP beads. Library concentration was determined by Qubit dsDNA High Sensitivity Kit.

Captured libraries were sequenced on MGISEQ-2000 sequencer (MGI, China) using the 2 × 100 paired-end sequencing method per manufacturer’s instruction.

### Preparation of two-human cfDNA blend sample

White blood cells from the two donors were first sequenced to determine genotypes. 11 heterozygous from the “spike-in” donor and 58 homozygous single nucleotide polymorphisms (SNPs) shared by the two donors covered by the IDT target capture panel were then selected to measure the sensitivity and specificity of variant detection, respectively. cfDNA samples of the two donors were mixed at a ratio of 1:200 to simulate cfDNA with a 0.25% “spike-in” variant allele frequencies (VAFs) using the heterozygous SNPs from the “spike-in” donor. Experiments were performed in duplicates.

### Data analysis

Adapter trimming and quality control of sequencing data were performed using Fastp (v0.19.7) [30]. Paired-end reads of targeted sequencing and targeted methylation sequencing were aligned to the hg19 reference human genome using bwa (v0.7.17) and BitMapperBS (v1.0.0.8), respectively [31, 32]. Duplications were marked and reads were deduplicated using sambamba (v0.6.8) [33]. Removal of sequencing errors using duplex UMIs and variant calling were performed using custom python scripts. Methylation rates of cytosines were calculated as #C/(#C+#T) for each CpG site with at least 4x coverage, and M-bias of sequence reads was analyzed using MethylDackel (v0.3.0) (https://github.com/dpryan79/MethylDackel). The cytosine conversion rate was calculated using the methylation ratio of the spiked-in lambda DNA. GC-bias metrics were analyzed using Picard Tools (v 2.10.10) (http://broadinstitute.github.io/picard). Insert size distribution, base distribution of reads, on-target rate, and sequencing depth were analyzed using custom Perl scripts.

### Data Access

The data that support the findings of this study have been deposited in the CNSA (https://db.cngb.org/cnsa/) of CNGBdb with accession number CNP0001331.

## Results

### Comparison of dsLib *vs*. ssLib method for targeted deep sequencing of cfDNA

We first compared double-stranded library (dsLib) preparation and single-stranded library (ssLib) preparation methods for cfDNA mutation detection using deep sequencing (Figure 1A, see Methods for more details). KAPA Hyper Prep kit, a widely used NGS library construction kit which is based on the conventional dsDNA library preparation methodology, was also included as a reference to evaluate performance of our self-developed dsLib workflow. Duplex unique molecular identifier (UMI)-based adapters were used to reduce noises that may derive from PCR and/or sequencing errors [21] (see Methods for more details). Since in clinical practice the amount of extracted cfDNA was often limited and highly variable [34], we used 1 ng, 5 ng, and 10 ng cfDNA as inputs for library construction respectively (Supplementary Table 1). Prepared libraries underwent hybridization-based target enrichment procedure and captured libraries were sequenced to > 20000x raw average depth (see Methods for more details). Results showed that library yields were similar between dsLib and ssLib workflow (Supplementary Figure 1A). The two workflows also achieved similar deduplicated depths (Figure 1B and Supplementary Table 1). Yet, the ssLib workflow was more complicated and time-consuming than the dsLib method (8h vs 3.5h, see Methods for more details). Remarkably, our self-developed dsLib protocol showed significantly better performance than the commercial KAPA workflow (Figure 1B and Supplementary Figure 1).

**Figure 1:**
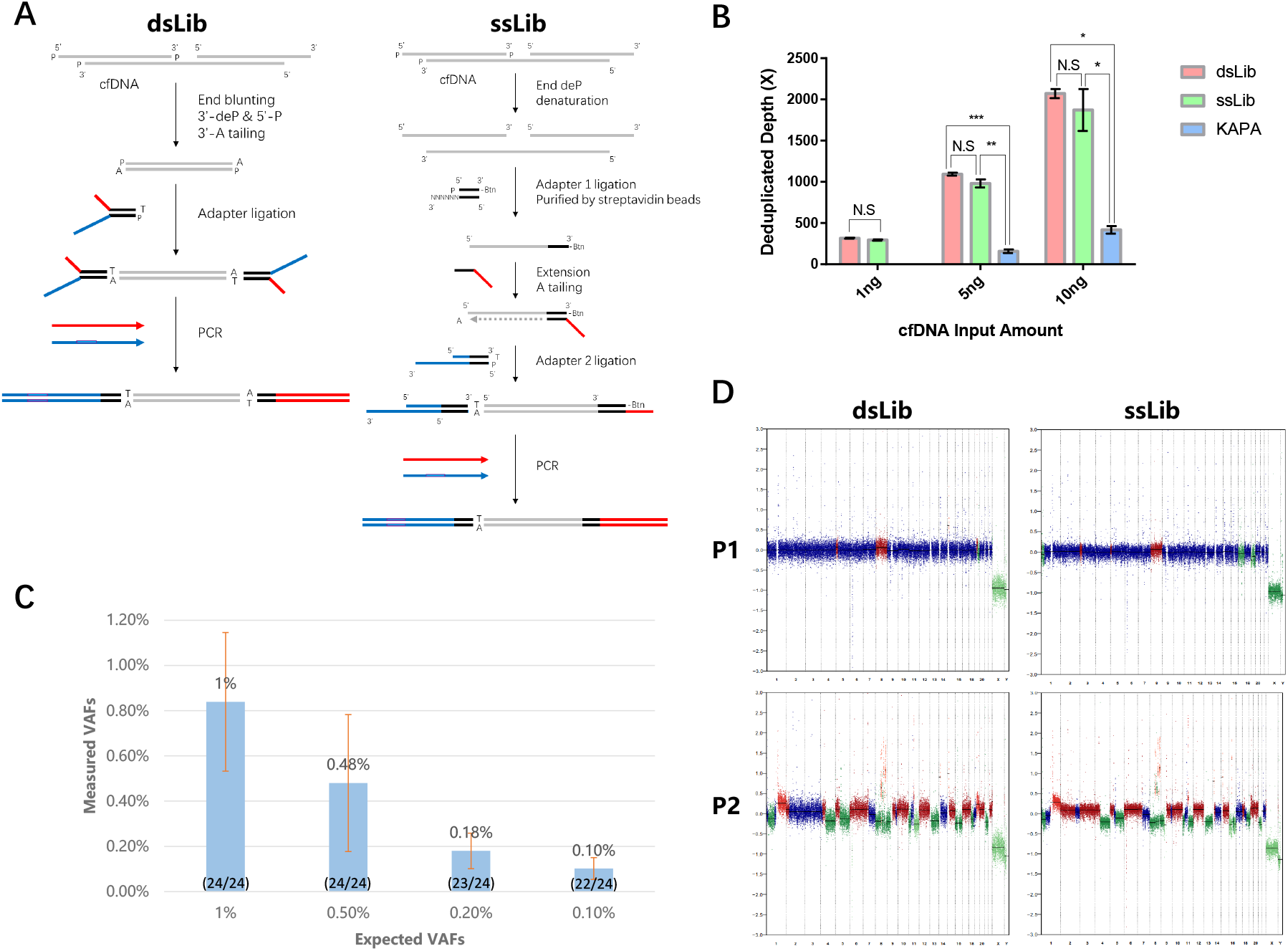
Comparison of dsLib and ssLib workflow for cfDNA mutation detection by targeted deep sequencing. **(A)** Schematic view of our self-developed dsLib and ssLib workflow. See Methods for more details. **(B)** Deduplicated depths of libraries constructed by dsLib, ssLib, and the KAPA kit. Duplicates were performed for each experimental condition. Data are presented as mean ± SD. N.S, p>=0.05; *, 0.01<=p<0.05; **, 0.001<=p<0.01; ****, p<0.0001, as calculated by Student’s t-test. **(C)** Detection of low VAF mutations by dsLib workflow in simulated cfDNA samples. Triplicates were performed for each experimental condition. Data are presented as mean ± SD. The numbers in parentheses represent the number of detected mutations/total mutations. **(D)** CNVs detected by dsLib and ssLib methods respectively, in plasma cfDNA samples from lung cancer patients P1 and P2. X-axis, chromosome. Y-axis, CNV adjusted by GC content and map-ability.

To further validate its ability to detect low abundance mutations in cfDNA and confirm the limitation of detection (LOD), we applied our dsLib workflow on 40 ng cfDNA spiked-in with cfDNA reference standards, simulating cfDNA samples with defined variant allele frequencies (VAFs) (0.1%, 0.25%, and 0.5%, see Methods for more details). 100% (24/24), 100% (24/24), 95.8% (23/24), and 91.7% (22/24) mutations were detected in cfDNA samples with 1%, 0.5%, 0.25% and 0.1% expected VAFs respectively, showing good correlation between the measured and expected VAFs (Figure 1C). The analytical performance of our assay was also evaluated using two-human cfDNA blend samples (see Methods for more details) to more closely mimic cfDNA carrying low VAF mutations. Briefly, single nucleotide polymorphism (SNP) sites where the “spike-in” donor carries heterozygous alleles while the “background” donor carries homozygous alleles were used to evaluate assay sensitivity; SNP sites where the “spike-in” donor and “background” donor carry the same homozygous alleles were used to evaluate assay specificity. We obtained a sensitivity of 95.5% (21/22 SNPs evaluated) and a specificity of 99.1% (115/116 SNPs evaluated) using the UMI error correction. Sensitivity was slightly lower (86.4%; 19/22 SNPs evaluated) if only variants supported by at least one duplex UMI family are considered true variants, while specificity was further improved to 100% (116/116) (Supplementary Table 2). The results indicated that our custom dsLib workflow provides satisfactory sensitivity for detection of low abundance variants in cfDNA.

ctDNA has been proven to be shorter than cfDNA originated from normal cells [35, 36]. Theoretically, the ssLib workflow preferentially enriches short DNA molecules and therefore may enrich ctDNA and improve its detection [28]. Copy number variation (CNV) is a hallmark of cancer and could be used as a biomarker for ctDNA [37]. Here, we compared CNV detectability of plasma cfDNA from lung cancer patients using either dsLib or ssLib workflow to test the hypothesis that ssLib may enrich for shorter ctDNA. We found no significant difference in CNV detection by ssLib workflow *vs*. dsLib (Figure 1D), consistent with previous study which showed that ssDNA-based workflow did not enrich for fetal DNA for NIPT, despite the finding that it did enrich for shorter cfDNA fragments [38]. Taken together, our results suggest that dsLib workflow is more preferable for ctDNA mutation detection.

### Comparison of dsLib *vs*. ssLib for cfDNA methylation sequencing

Bisulfite sequencing has been a widely used sequencing technology for methylation profiling, where methylation status of cytosines could be determined at single-nucleotide resolution. This technology leverages the fact that methylated cytosine remains unaffected when treated with sodium bisulfite, whereas unmethylated cytosine is converted to uracil [39].

To compare performance of the single-stranded methylation sequencing library construction (ssmLib) and the double-stranded methylation sequencing library construction (dsmLib) (Figure 2A), we applied these two workflows on 1 ng, 5 ng, and 10 ng cfDNA as inputs and captured the libraries with a 198 kb methylation capture panel (Supplementary Table 3). Sequencing results showed that ssmLib produced significantly higher library yields and deduplicated depths than dsmLib; the on-target rates were also slightly higher in ssmLib libraries than dsmLib libraries (Figure 2 B-C and Supplementary Figure 2). Notably, libraries produced by ssmLib had more short insert fragments than those produced by dsmLib (Figure 2D). These results can be attributed to DNA degradation caused by the bisulfite conversion process, which involves high temperature and low pH conditions [40]: during ssmLib workflow, the resulted short cfDNA fragments can still be captured by the ssDNA-based adapter ligation; on the other hand, during dsmLib workflow, since bisulfite was applied to the adapter-ligated dsDNA, excessive damage of the templates will cause the libraries to lack paired adapters and lost during subsequent amplification, resulting in much lower library yields and effective sequencing depths. For measurements of CpG site methylation level, technical replicates showed good correlation for both methods (Supplementary Figure 3) with various DNA input amounts (Figure 2E).

**Figure 2:**
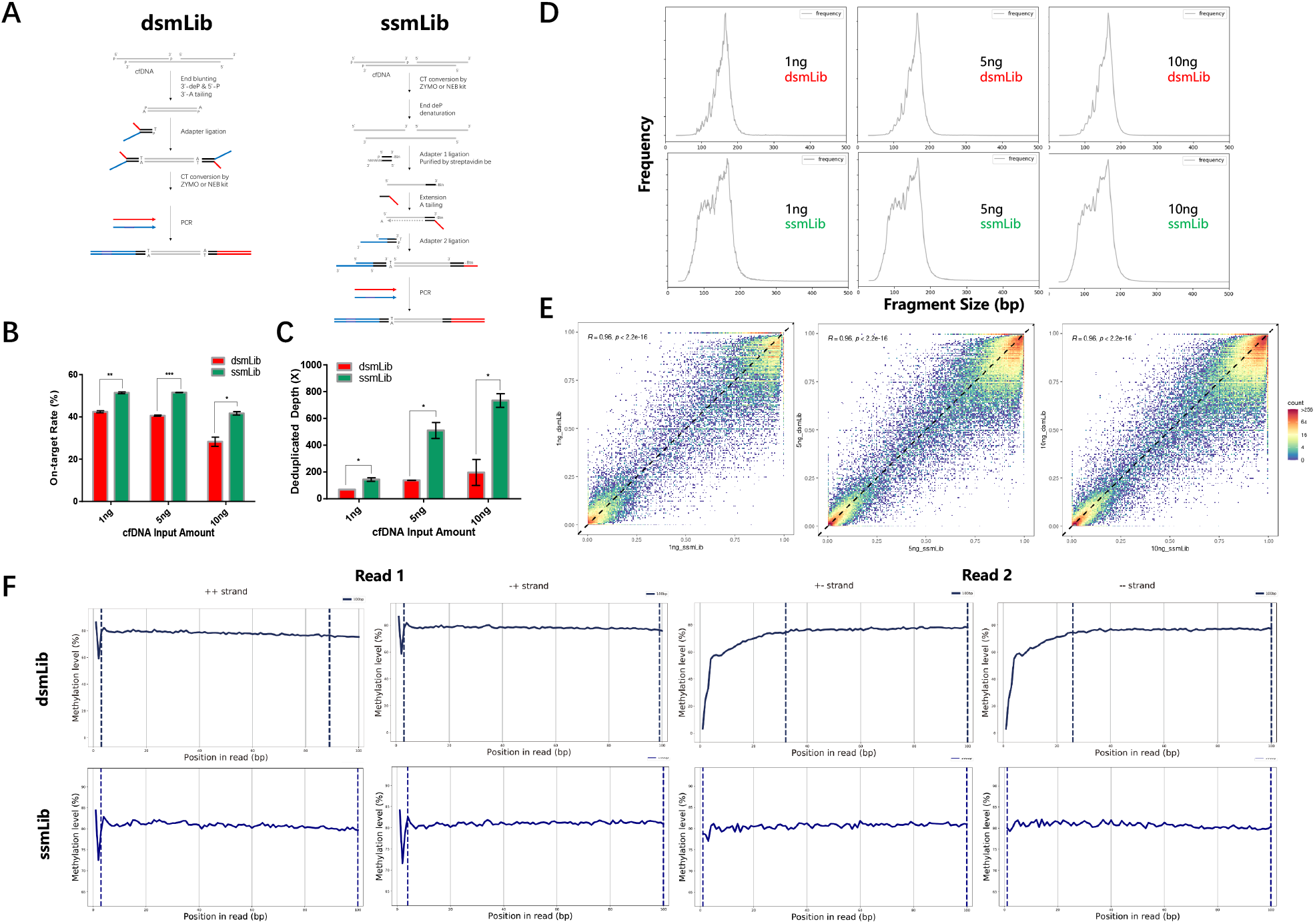
Comparison of dsmLib and ssmLib workflow for cfDNA methylation profiling by bisulfite sequencing. **(A)** Schematic view of our dsmLib and ssmLib procedures. **(B)** On-target rates and **(C)** deduplicated depths of libraries prepared by dsmLib and ssmLib workflow. Duplicates were performed for each experimental condition. Data are presented as mean ± SD. N.S, p>=0.05; *, 0.01<=p<0.05; **, 0.001<=p<0.01; ****, p<0.0001, as calculated by Student’s t-test. **(D)** Size distributions of library insert fragments. **(E)** Pearson correlation of methylation levels between ssmLib (x-axis) and dsmLib (y-axis) libraries. **(F)** M-bias plots of libraries prepared by dsmLib and ssmLib workflow. For each row from left to right: Read 1 ++ strand, Read 1 -+strand, Read 2 +-strand, and Read 2 --strand. X-axis, position in read (bp). Y-axis, methylation level (%).

Methylation bias (M-bias) is the term describing measured methylation levels that deviate from true values, often observed near the 3’ end of sequenced fragments due to unmethylated cytosines introduced by the end-repair step during dsDNA-based library preparation [41, 42]. Theoretically, libraries produced by ssmLib may show less to no M-bias since there is no end-repair step involved (Figure 2A). Indeed, we observed severe M-bias in Read 2 of dsmLib libraries, but not in ssmLib libraries (Figure 2F). Taken together, these results suggest that ssmLib method is more preferrable for the application of cfDNA methylation sequencing.

Recently, several enzyme-based cytosine conversion methods have been developed as gentler substitutes for bisulfite conversion [43, 44]. We also compared performance of a novel enzyme-based workflow (the NEB EM-seq Conversion Module) with the conventional bisulfite conversion workflow (using the widely used ZYMO EZ DNA Methylation Gold kit) (Figure 3A). EM-seq conversion module uses a two-step enzymatic conversion process to detect modified cytosines: the first step uses TET2 and an oxidation enhancer to protect modified cytosines from downstream deamination while converting 5-methylcytosine (5mC) to 5-carboxycytosine (5caC). The second step uses APOBEC to enzymatically deaminate cytosine but does not convert 5caC (the original 5mC). As expected, the ssmLib libraries produced by bisulfite conversion had more short insert fragments than those produced by enzyme-based conversion. Meanwhile, with ssmLib workflow, the bisulfite conversion method generated significantly higher library yields and deduplicated depths than enzymatic conversion, while similar cytosine conversion efficiencies were observed for the two methods (Figure 3C-E and Supplementary Table 3). For dsmLib workflow, however, there was no significant difference in either library yields, deduplicated depths, or fragment size distributions between the two conversion methods (Figure 3B, Supplementary Figure 4 and Supplementary Table 3). Among all, bisulfite conversion coupled with ssmLib workflow still achieved the highest deduplicated sequencing depth. Also, the enzymatic conversion is more time-consuming (8h vs 3.5h) than the bisulfite conversion. Taken together, our results favor bisulfite conversion coupled with ssLib workflow for cfDNA methylation sequencing.

**Figure 3:**
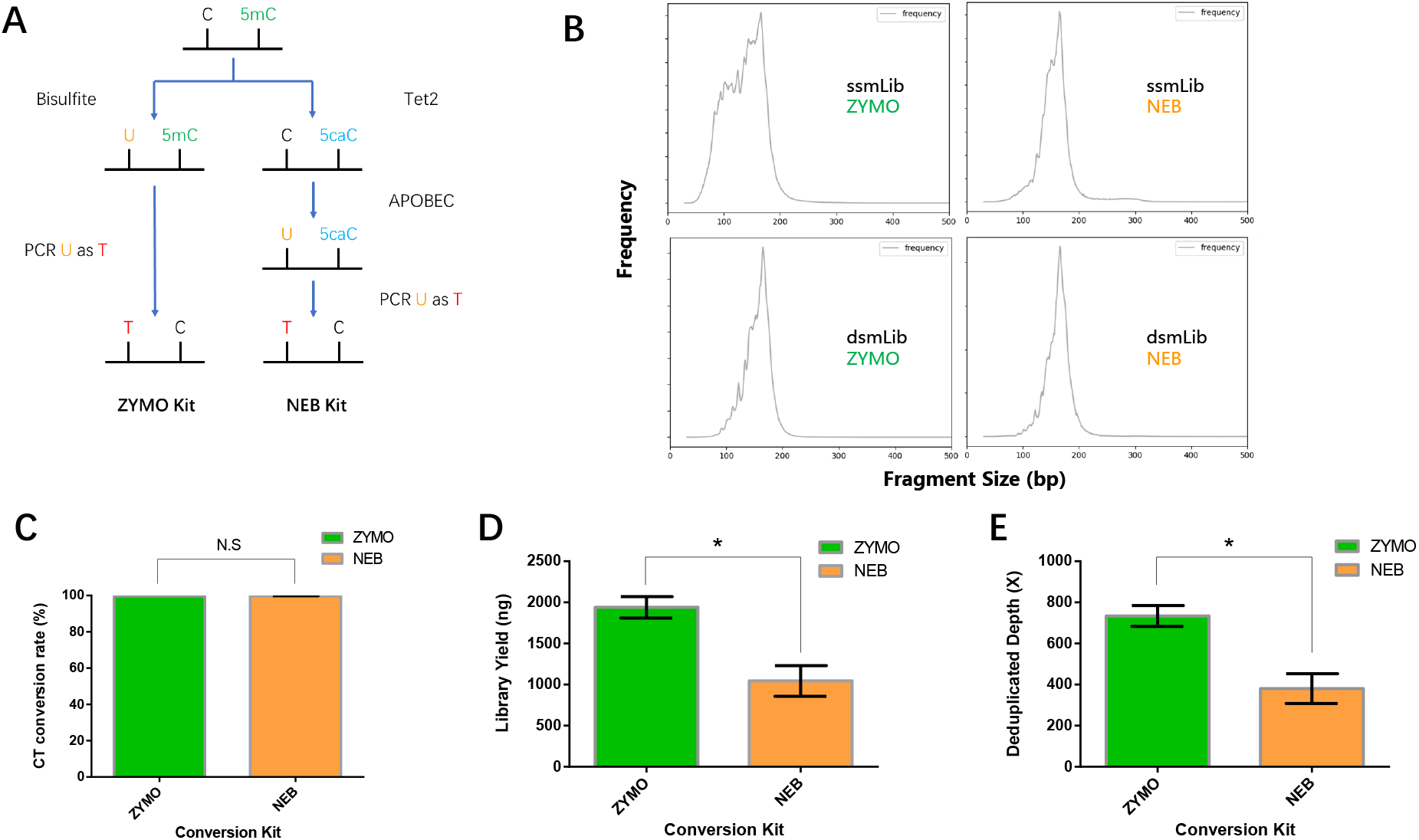
Comparison of the chemical and enzymatic cytosine conversion for cfDNA methylation sequencing. **(A)** Technical principles of bisulfite conversion and enzymatic conversion. 5caC, 5-carboxylcytosine. T, thymine. **(B)** Size distribution of library insert fragments. X-axis, fragment size (bp). Y-axis, frequency count. **(C)** CT conversion rates, **(D)** library yields, and **(E)** deduplicated depths of ssmLib libraries. Duplicates were performed for each experimental condition. Data are presented as mean ± SD. N.S, p>=0.05; *, 0.01<=p<0.05, as calculated by Student’s t-test.

## Conclusion

The double-stranded library preparation method is more advantageous for ctDNA mutation detection thanks to the higher data quality and easy workflow. Meanwhile, bisulfite conversion coupled with single-stranded library preparation showed overall better performance for cfDNA methylation sequencing. Our results suggest that when performing high-throughput sequencing for cfDNA, depending on the downstream applications, these two library preparation methods should be chosen accordingly.

## Discussion

In recent decades, thanks to the development of NGS technology, the cost of high-throughput DNA sequencing had dropped dramatically, making it affordable for researchers worldwide [45]. Library construction is a key step for successful NGS workflow and high-quality data generation. In this study, we compared dsDNA and ssDNA-based library construction methods for cfDNA deep sequencing (i.e., for ctDNA variant detection) and methylation profiling.

A major difference between dsDNA and ssDNA based cfDNA library construction methods is that cfDNA molecules harboring single-strand breaks (also called nicks) as well as those existing as ssDNA form could be utilized by the ssLib (or ssmLib) workflow but would not be ligatable when using the dsLib (or dsmLib) workflow (Figure 1A and 2A). Naturally nicked and/or single-stranded cfDNA molecules may only be a very small fraction hence this difference would be expected to be small and may not cause significant impact on the effective sequencing depth. Indeed, we observed similar deduplicated depth for cfDNA libraries generated using ssLib or dsLib workflow (Supplementary Figure 1B); in fact, deduplicated depth of ssLib libraries were even slightly inferior than dsLib, possibly due to the fact that ssLib workflow is lengthier and requires more beads purification and therefore may cause template loss.

Using detected CNV level as an indicator of ctDNA fraction, we also showed that there was no significant enrichment of ctDNA by ssLib compared to dsLib workflow (Figure 1D), consist with previous research conducted in the setting of NIPT which showed that ssLib workflow does not enrich for shorter fetal DNA [28, 38]. It was suggested that intrinsic biological differences between fetal DNA and maternal DNA molecules might account for the failure of ssDNA workflow to enrich for fetal DNA [28, 38], and similar mechanism may also explain our results for ctDNA. Further study is needed to deepen our understanding of cfDNA/ctDNA generation processes and/or to develop novel library construction methods for ctDNA enrichment.

Importantly, application of dsLib workflow further allows utilization of duplex UMIs, which make it possible to recover original dsDNA fragments following paired-end sequencing and utilize the information from complementary strands of DNA molecules to correct possible PCR and/or sequencing errors, achieving an extra low base error rate and higher specificity with variant detection [21]. Taken together, our results demonstrate that current state-of-the-art dsDNA-based library preparation is more preferable for the application of deep sequencing for ctDNA variant detection.

On the contrary, a clear advantage was observed for ssmLib libraries for bisulfite sequencing compared to dsmLib (Figure 2B-C). This is because libraries were constructed before bisulfite conversion during the dsmLib workflow (Figure 2A), and the nicked DNA resulting from the bisulfite conversion won’t be sequenced due to the lack of paired adapters. During the ssmLib workflow, however, cfDNA ligation happens after the bisulfite treatment, where the nicked and single-stranded DNA molecules resulting from the bisulfite treatment can still be ligated with adapters, therefore preserving more DNA templates for sequencing (Figure 2A and 2D), eventually achieving a higher effective depth.

Theoretically, gentler enzyme-based cytosine conversion method would avoid the assumed template loss caused by bisulfite treatment on the adapter-ligated library fragments and may therefore greatly improve the results of dsLib workflow when used for methylation profiling. Our results, however, still favored the bisulfite conversion for both dsmLib and ssmLib workflow due to the higher library yields as well as higher deduplicated depths, suggesting that there may be excessive loss of templates during the enzyme-conversion workflow (Figure 3C-E and Supplementary Figure 4). Indeed, this may be attributed to the two rounds of beads purification in the enzyme-based conversion. Also, the current enzyme-based conversion workflow is more labor- and time-consuming compared to the bisulfite conversion. Development of more effective enzyme-based cytosine conversion methods may require improvements in template recovery and further simplification of the workflow.

In addition, methylation bias (M-bias) was proposed to be an important library preparation quality metric for methylation profiling, since its existence could cause significant bias in measurements of methylation level [41, 42]. M-bias is caused by the end-repair step in the conventional dsmLib workflow which typically recruits unmethylated cytosines instead of methylated cytosines during the fill-in reaction (Figure 2A). The filled-in cytosines were then converted to uracils regardless of the original cytosine methylation status in the genome, resulting in incorrect methylation level being assigned to the 3’ end of the sequenced reads [41, 42]. The ssmLib method could perfectly overcome this problem since it does not involve an end-repair step and is a post–bisulfite conversion library construction method (Figure 2A and 2F), adding another advantage to the ssmLib method. Collectively, our results favor the use of ssmLib workflow for cfDNA methylation profiling. Our findings could help researchers maximize the efficiency of NGS library preparation and produce better quality sequencing data.

---

**Supplementary Figure 1: (A) Library yields and (B) mean fragment lengths of sequenced libraries constructed by dsLib, ssLib, and the KAPA workflow**.

Duplicates were performed for each experimental condition. Data are presented as mean ± SD. N.S, p>=0.05; *, 0.01<=p<0.05; **, 0.001<=p<0.01; ****, p<0.0001, as calculated by Student’s t-test.

**Supplementary Figure 2: Library yields by dsmLib and ssmLib**. Duplicates were performed for each experimental condition. Data are presented as mean ± SD. N.S, p>=0.05; *, 0.01<=p<0.05; **, 0.001<=p<0.01, as calculated by Student’s t-test.

**Supplementary Figure 3: Pearson correlation of methylation levels between (A) dsmLib or (B) ssmLib libraries**.

**Supplementary Figure 4: (A) CT conversion rates, (B) library yields, and (C) deduplicated depths of dsmLib libraries using bisulfite and enzymatic conversion**. Duplicates were performed for each experimental condition. Data are presented as mean ± SD. N.S, p>=0.05; *, 0.01<=p<0.05, as calculated by Student’s t-test.

