## Supplementary figures and images for "Comparison of dsDNA and ssDNA-based NGS library construction methods for targeted genome and methylation profiling of cfDNA"

### Supplementary Figure 1

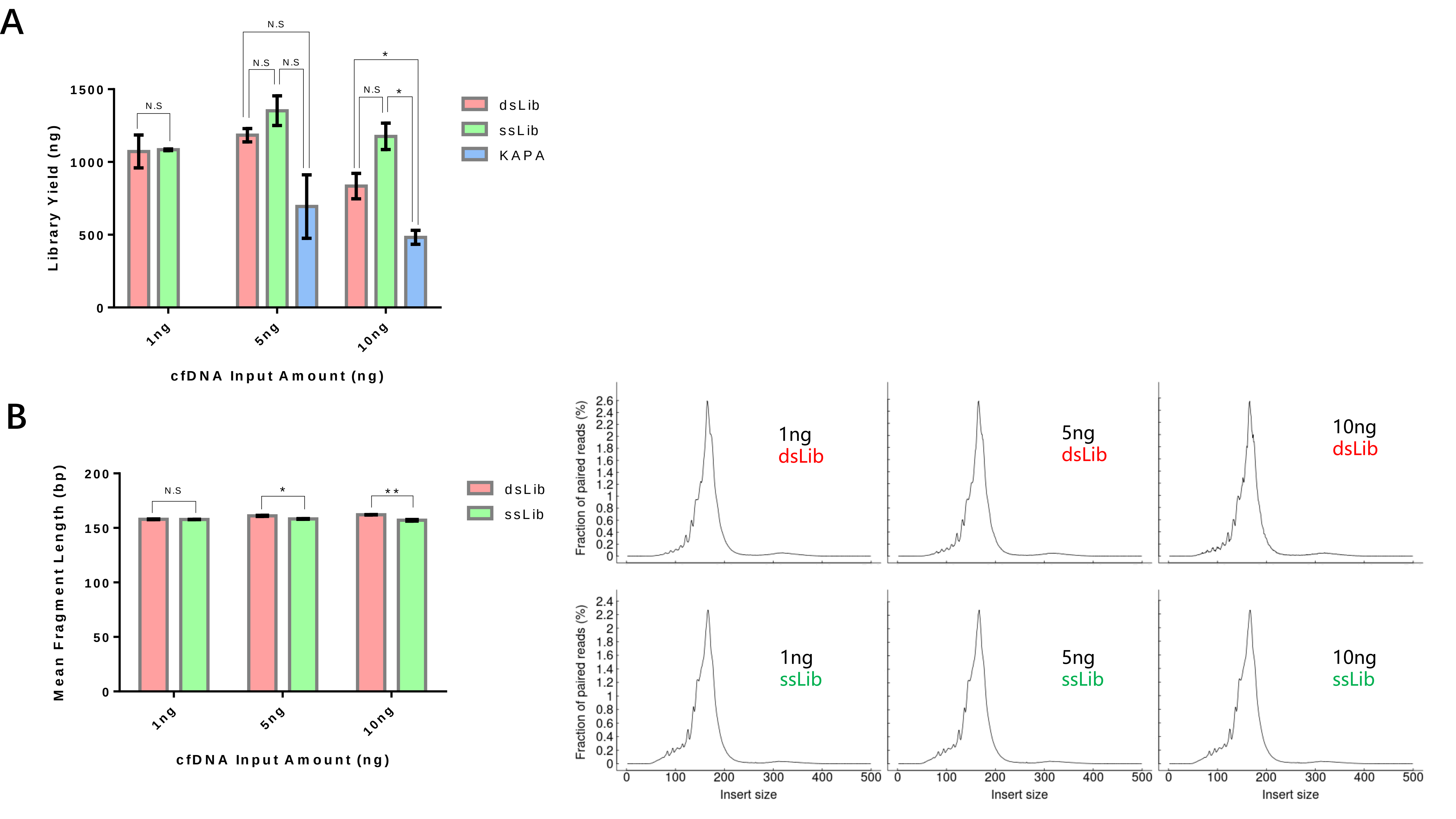

### Supplementary Figure 2

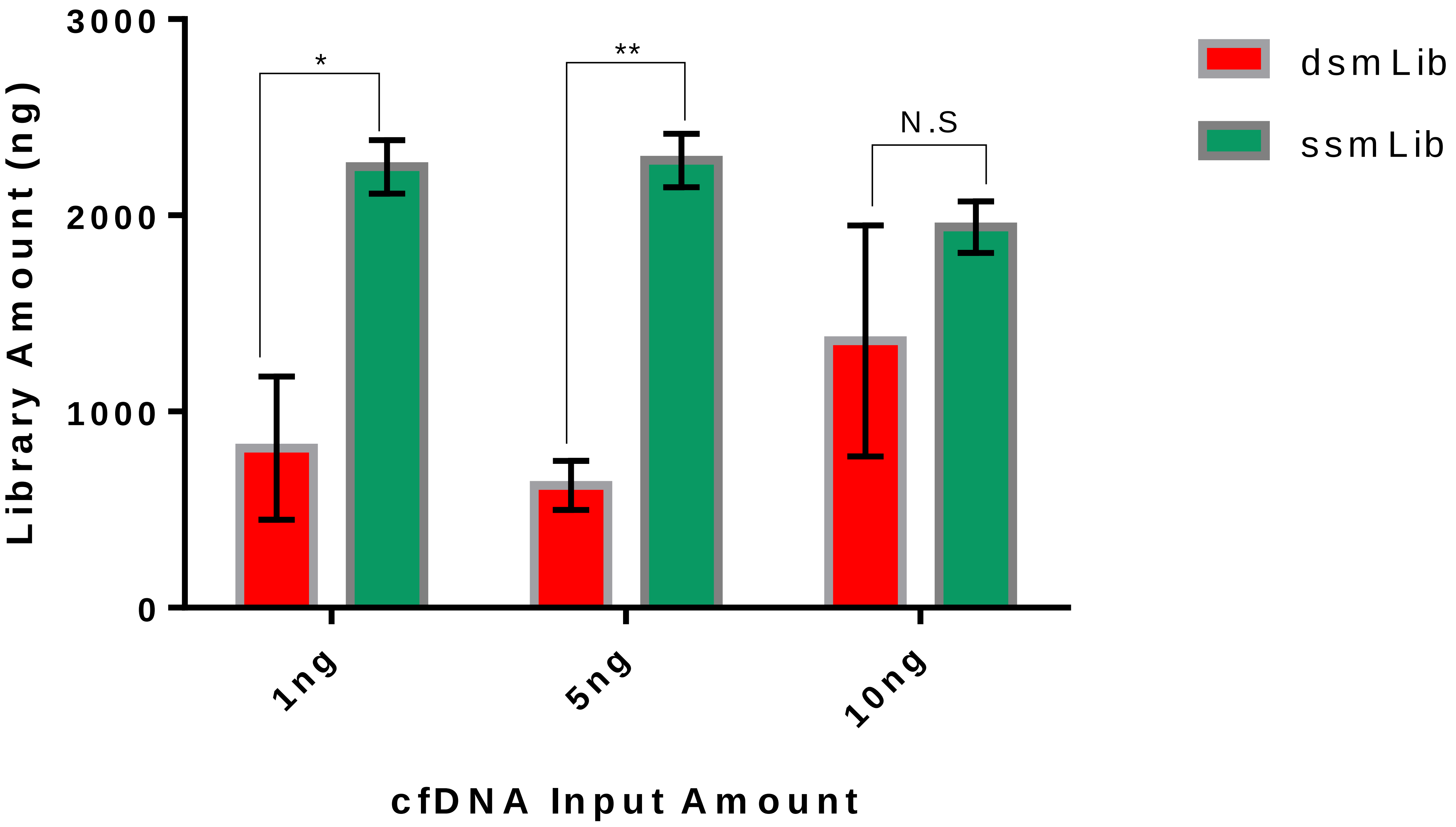

### Supplementary Figure 3

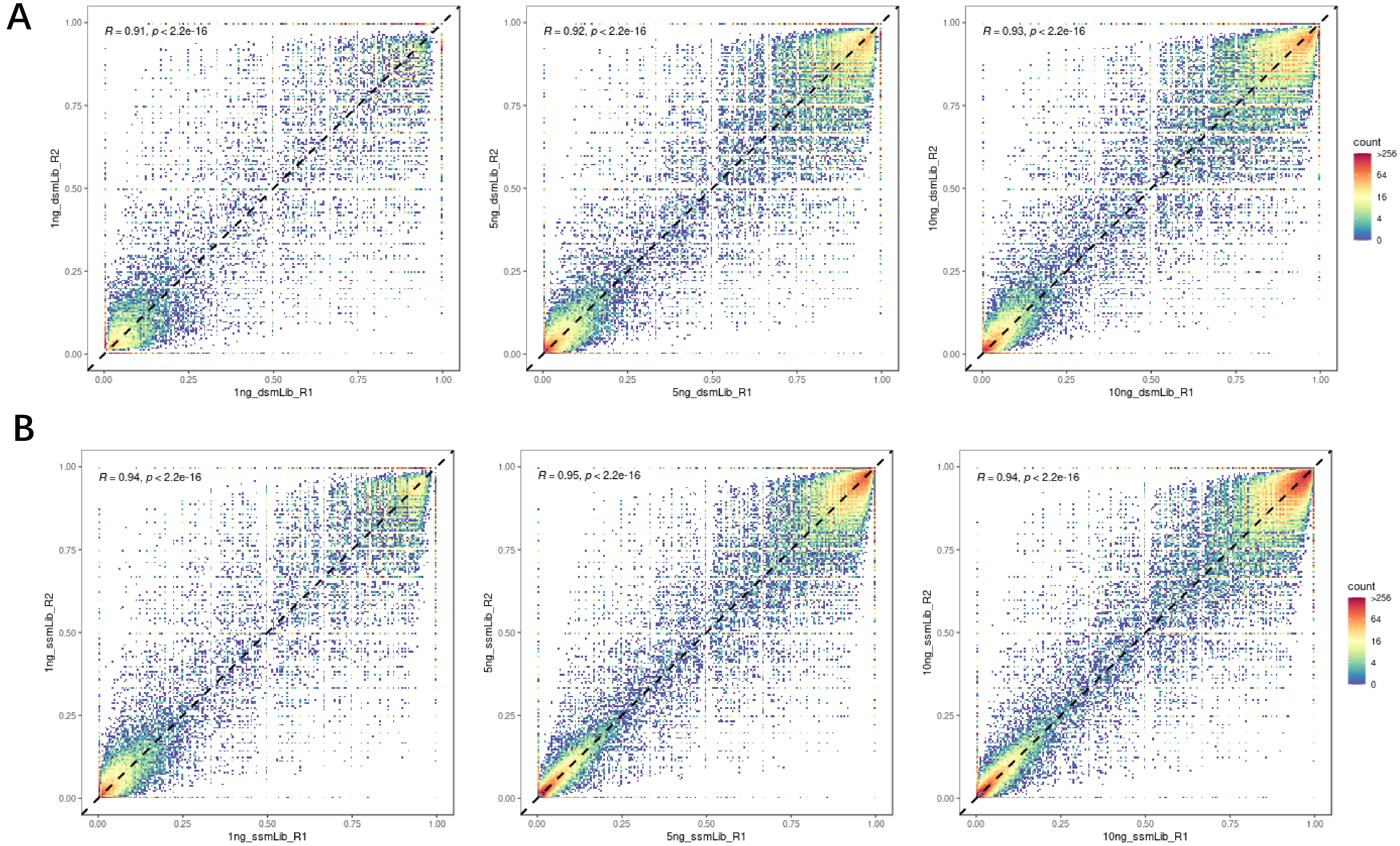

### Supplementary Figure 4

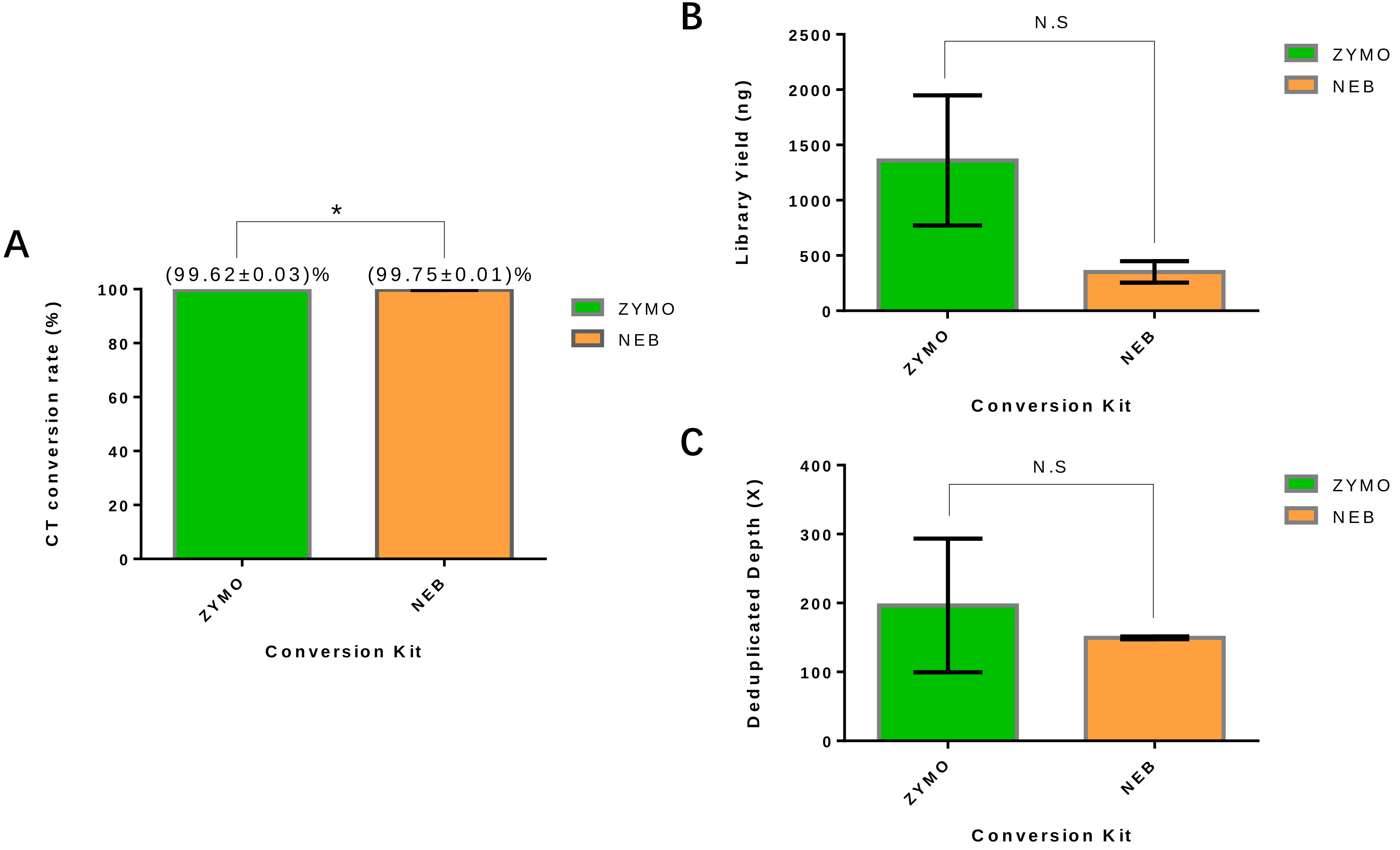
